# BigBrainWarp: Toolbox for integration of BigBrain 3D histology with multimodal neuroimaging

**DOI:** 10.1101/2021.05.04.442563

**Authors:** Casey Paquola, Jessica Royer, Lindsay B. Lewis, Claude Lepage, Tristan Glatard, Konrad Wagstyl, Jordan DeKraker, Paule-J Toussaint, Sofie L Valk, Louis Collins, Ali R. Khan, Katrin Amunts, Alan C. Evans, Timo Dickscheid, Boris Bernhardt

## Abstract

Neuroimaging stands to benefit from emerging ultrahigh-resolution histological atlases of the human brain; the first of which is “BigBrain”. Ongoing research aims to characterise regional differentiation of cytoarchitecture with BigBrain and to optimise registration of BigBrain with standard neuroimaging templates. Together, this work paves the way for multi-scale investigations of brain organisation. However, working with BigBrain can present new challenges for neuroimagers, including dealing with cellular resolution neuroanatomy and complex transformation procedures. To simplify workflows and support adoption of best practices, we developed BigBrainWarp, a toolbox for integration of BigBrain with multimodal neuroimaging. The primary BigBrainWarp function wraps multiple state-of-the-art deformation matrices into one line of code, allowing users to easily map data between BigBrain and standard MRI spaces. Additionally, the toolbox contains ready-to-use cytoarchitectural features to improve accessibility of histological information. The present article discusses recent contributions to BigBrain-MRI integration and demonstrates the utility of BigBrainWarp for further investigations.

## 1. Introduction

Understanding brain anatomy benefits from a multi-scale perspective, from the microscopic to the macroscopic level. Regional variations in cells underlie macro-scale patterns, whether they are reflective of functional dynamics, age, or disease states. For over 150 years (von Gudden, 1886), histological analysis of *post mortem* tissue has helped to reveal the microscopic architecture of the brain. Neuroanatomists observed a distinctive layered organisation of cells within the cortex (Baillarger, 1840) and developed principles of cortical organisation, including the definition of cortical types (Meynert, 1867), cortical areas (Brodmann, 1908; Geyer et al., 2011; Von Economo and Koskinas, 1925), and cortical gradients (Bailey and von Bonin, 1951; Goulas et al., 2019; Sanides, 1962). More recently, digitisation of *post mortem* tissue has allowed automated characterisation of cytoarchitecture (Schleicher et al., 1999). This mergence of histology with computational neuroscience supports more observer-independent evaluation of classic principles (Amunts et al., 2020; Paquola et al., 2019; Schiffer et al., 2020; Spitzer et al., 2018) and paves the way for novel investigations of the cellular landscape of the brain.

*In vivo* neuroimaging offers a complementary window into the functional dynamics of the brain. Additionally, the non-invasive nature of magnetic resonance imaging (MRI) supports examination of population-level variation, which is largely inaccessible to *post mortem* neuroanatomy. Human brain mapping research has furthermore established standard spaces, notably the MNI152 space for volumetric whole-brain analysis (Fonov et al., 2011b, 2009; Mazziotta et al., 2001a, 2001b) and “fsaverage” and “fs_LR” for surface-based cortical analyses (Fischl et al., 1999; Van Essen et al., 2012). Despite ongoing advances in attaining higher spatial resolution with higher field strength (Deistung et al., 2013; Holdsworth et al., 2019; Sitek et al., 2019; Trampel et al., 2019; Turner and De Haan, 2017), *in vivo* MRI researchers remain constrained by limited spatial resolution from making inferences on a cellular level. Establishing the relation between macro-scale patterns and cellular architecture is crucial to substantiate physiological patterns observed with MRI and for further development of brain-inspired computational models.

BigBrain is a singular 3D volumetric reconstruction of a sliced and cell-body stained complete human brain (Amunts et al., 2013). This resource allows for computational analysis of an entire human brain in relation to cell staining at high resolutions (up to 20μm). Tailored for neuroimagers, it is available in common MRI formats (minc and NifTI), accompanied by cortical surface reconstructions (Lewis et al., 2014), and nonlinearly registered to standard MRI templates (ICBM152 and MNI-ADNI) (Fonov et al., 2011a). Furthermore, recent studies have expanded the resource by offering improved registrations to standard spaces (Lewis et al., 2020; Xiao et al., 2019), nuanced intracortical surface models and laminar approximations (Wagstyl et al., 2020, 2018a) as well as regional segmentations (DeKraker et al., 2019; Xiao et al., 2019). Several studies have already capitalised on this unique resource for integrative histological-neuroimaging analyses, including comparison of cytoarchitectural and functional gradients (Paquola et al., 2019), cross-validation of *in vivo* defined microstructural gradients in the insula with histological measures (Royer et al., 2020), mapping variations in functional connectivity along the histological axis of the mesiotemporal lobe (Paquola et al., 2020b), fMRI responses of the histologically-defined auditory system (Sitek et al., 2019), comparison of cytoarchitectural similarity with MRI-derived estimates of structural connectivity (Wei et al., 2018), evaluating the cytoarchitectural heterogeneity of the default mode network (Paquola et al., 2021), and analyses of the cytoarchitectural similarity of large-scale network hubs (Arnatkevičiūtė et al., 2020).

The present article introduces the BigBrainWarp toolbox. The aim of the toolbox is to facilitate integration of BigBrain with neuroimaging modalities, helping neuroscientists to utilise cytoarchitectural information in conjunction with *in vivo* imaging. The toolbox is open and includes (i) histological features and pre-transformed maps in BigBrain and imaging spaces, (ii) codes for performing data transformations and (iii) a knowledgebase for multi-modal integration of BigBrain with MRI. Toolbox functions and tutorials are documented on http://bigbrainwarp.readthedocs.io. Here, introduce BigBrain to new users and demonstrate the utility of the BigBrainWarp toolbox. In Section 2, we overview the derivation of cytoarchitectural features from BigBrain and survey recent contributions to BigBrain-MRI integration. These include publication of histological cortical maps, regional segmentations, and registration efforts. Finally, we detail the core functions of BigBrainWarp and the current contents of the toolbox. In Section 3, we share three tutorials to illustrate potential applications of BigBrainWarp.

## 2. Material and Methods

### Overview of BigBrain

In brief, the reconstruction of BigBrain involved coronal slicing of a complete paraffin-embedded brain (65-year-old male) into 7400 sections at 20μm thickness. Each section was stained for cell bodies (Merker, 1983), digitised and subjected to manual and automatic artefact repair. The digitised sections were reconstructed into a contiguous 3D volume. The volumetric reconstruction is available online at 40μm, 100μm, 200μm, 300μm, 400μm and 1000μm resolutions (http://bigbrainproject.org). The 40μm version is released as 125 individual blocks corresponding to five subdivisions in the *x, y*, and *z* directions, with overlap. 100-1000 μm resolutions are provided as single files. Merker staining used in BigBrain is a form of silver impregnation for cell bodies that produces a high contrast of black pigment in cells on a virtually colorless background (Merker, 1983). In the digitised images, darker colouring is represented by lower numbers (8bit graphics: 0-2^8^=black-white). It is common practice to invert the values of the intensity, such that image intensity increases with staining intensity.

The grey and white matter boundaries of the cortical surface released in 2014 contain 163,842 vertices on each hemisphere, with vertices aligned between pial and white surfaces (Lewis et al., 2014). Surfaces were generated using a modified version of CIVET (Kim et al., 2005; MacDonald et al., 2000). Since then, a number of additional surface reconstructions have been published from which we may attain a range of metrics (**Table 1**).

**Table 1:**
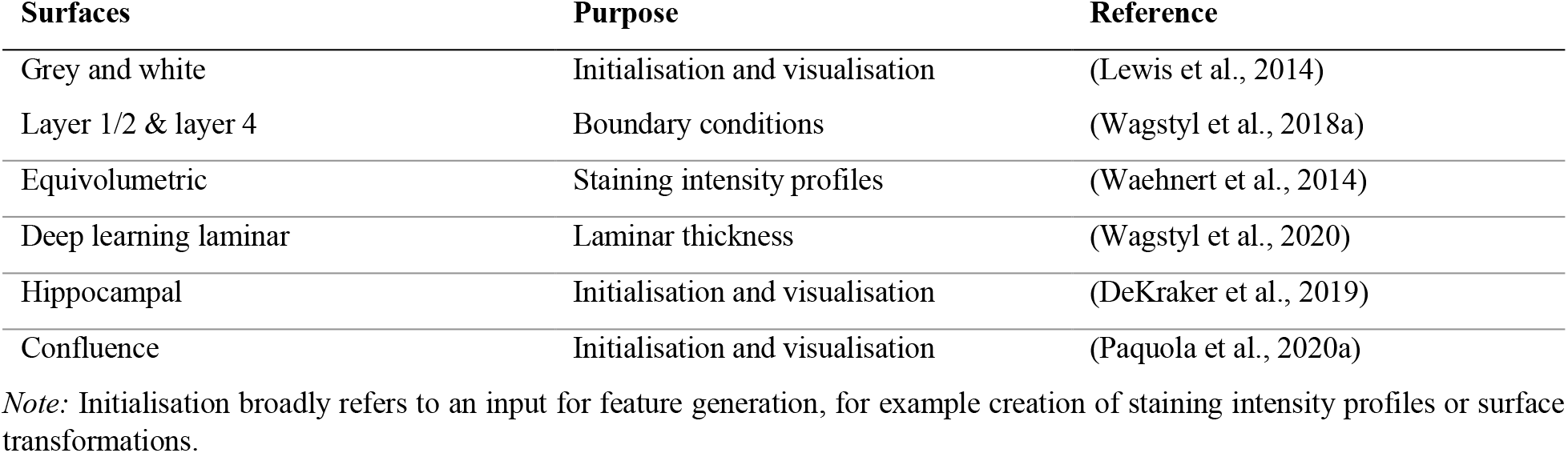
Surface constructions for BigBrain.

*Note:* Initialisation broadly refers to an input for feature generation, for example creation of staining intensity profiles or surface transformations.

### Staining intensity profiles and derived features

Sampling staining intensity from many cortical depths provides a profile of the cytoarchitecture, hereafter referred to as a staining intensity profile. This is achieved by constructing a set of surfaces within the cortex, then sampling intensity estimates at matched vertices across the surfaces. The current approach involves equivolumetric surface construction, whereby a set of intracortical surfaces are initialised at equidistant depths, then modulated by cortical curvature (Waehnert et al., 2014). This holds advantages for histological data because laminae vary in thickness depending on cortical folding (Bok, 1929). The procedure can be deployed using dedicated python scripts (Wagstyl et al., 2018b) and is implemented in BigBrainWarp.

Smoothing can be employed in tangential and axial directions to ameliorate the effects of artefacts, blood vessels, and individual neuronal arrangement (Wagstyl et al., 2018a). Smoothing across depths is enacted for each staining profile independently. Here, we use an iterative piece-wise linear procedure that minimises curve shrinkage, where the degree of smoothing is modulated by the number of iterations (Taubin, 1995). In contrast, surface-wise smoothing is performed at each depth and involves moving a Gaussian kernel across the surface mesh. We tested the impact of number of surfaces and smoothing on profiles, using the 100μm whole brain volume. Specifically, we evaluated spatial autocorrelation and number of profile peaks for each combination (number of surfaces 50-100, iterations of depth-wise smoothing=2-10, FWHM of surface-smoothing=0-8, **Figure S1**). Spatial autocorrelation was calculated as the average product-moment correlation of staining intensity profiles at various distances along the BigBrain surface mesh (distances: 1-50 steps). Increasing the number of surfaces beyond 50 did not impact the spatial autocorrelation and led to small increases in the number of peaks in intensity profiles. Depth-wise smoothing did not impact either outcome measure. As could be expected, surface-wise smoothing substantially increased spatial autocorrelation. For the initial BigBrainWarp release, we selected 50 surfaces, 2 iterations of depth-wise smoothing and (a modest) 2 FWHM surface-wise smoothing. BigBrainWarp also provides a simple function for generating staining intensity profiles.

Previous research has sought to characterise the laminar structure of the cortex using BigBrain staining intensity profiles (Paquola et al., 2019; Schleicher et al., 1999; Wagstyl et al., 2018a; Zilles et al., 2002). The isocortex generally contains six layers (Brodmann, 1909), certain features of which manifest on BigBrain staining intensity profiles. The transition from layer I to II exhibits a sharp increase in staining, because layer I is only sparsely populated with cells. Layer IV harbours a noticeable peak in cell staining, corresponding to dense packing of granule cells. The peak of layer IV corresponds to the division between supragranular and infragranular layers, which have markedly different roles in neural communication (Buffalo et al., 2011; Felleman and Van Essen, 1991; Rockland and Pandya, 1979). The relative depth of layer IV is also potentially informative, likely related to the propensity for feedforward vs feedback communication (Beul et al., 2017; Sanides, 1962; Wagstyl et al., 2018a), though the demarcation of feedforward and feedback projections is more multifactorial and complex (Rockland, 2015). A six-layered decomposition of BigBrain cortex has also been produced by training a convolutional neural network on manual annotations in 51 regions, then extending the model to the whole isocortex (Wagstyl et al., 2020) (**Figure 1E**). Laminar thickness estimates aligned with prior histological studies (Von Economo and Koskinas, 1925), while increasing overall spatial precision. There remains difficulty in extending these approaches to cortex without clear laminar differentiation, however (*i.e.,* anterior insula, mesiotemporal lobe).

**Figure 1:**
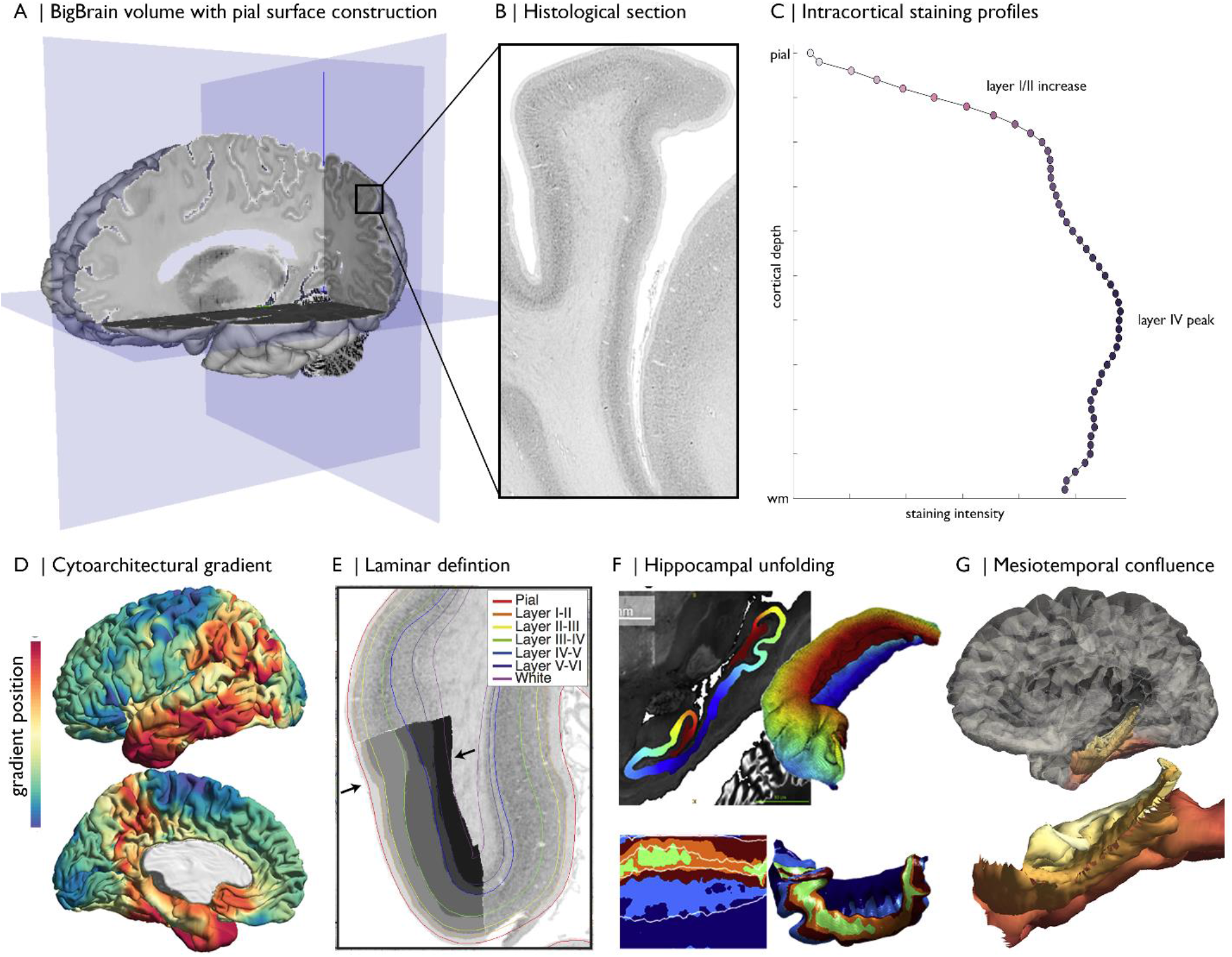
Magnification of cytoarchitecture using BigBrain, from **(A)** whole brain 3D reconstruction (taken on https://atlases.ebrains.eu/viewer) to **(B)** a histological section at 20μm resolution (available from bigbrainproject.org) to **(C)** an intracortical staining profile. The profile represents variations in cellular density and size across cortical depths. Distinctive features of laminar architecture are often observable i.e., a layer IV peak. Note, the presented profile was subjected to smoothing as described in the following section. BigBrainWarp also supports integration of previous research on BigBrain including **(D-E)** cytoarchitectural and **(F-G)** morphological models (DeKraker et al., 2019; Paquola et al., 2020a, 2019; Wagstyl et al., 2020).

More detailed characterisation of cytoarchitecture is offered by moment-based parameterisation of intracortical intensity profiles. This technique, pioneered by the Jülich group (Schleicher et al., 1999; Zilles et al., 2002), involves calculating the central moments *(i.e.,* mean, standard deviation, skewness, and kurtosis) of each staining intensity profile and the derivative profile, resulting in a multidimensional feature vector for each cortical point. Each central moment may be interpreted in neurobiological terms (Zilles et al., 2002). For example, mean intensity generally increases in the anterior to posterior direction and has been related to overall cellular density (Wree et al., 1982). In contrast, skewness varies from sensory to limbic areas (*i.e.,* sensory-fugal) and indexes the balance of cellular density in infra- vs supra-granular layers (Paquola et al., 2020b). Comparison of profiles can illuminate large-scale patterns of cortical organisation. Observer-independent discrimination of cortical areas can be accomplished by comparing moment-based feature vectors between neighbouring vertices (Schleicher et al., 1999). The areal boundaries are defined where the feature vector exhibits a sudden shift. Over the past 20 years, this procedure has been employed in 23 *post mortem* brains, including BigBrain, resulting in a 3D probabilistic atlas of the human brain (Amunts et al., 2020). While this work is based on a selection of histological sections of each brain, recent work investigates solutions for mapping each section in a stack with the help of deep learning, in order to produce gapless 3D maps at full detail (Schiffer et al., 2020) and ultimately obtain a dense mapping of the BigBrain model.

Cortex-wide cytoarchitectural similarity may also be estimated, by cross-correlating staining intensity profiles between different cortical locations (Paquola et al., 2019). We recently applied diffusion map embedding, a nonlinear manifold learning technique (Coifman and Lafon, 2006), to the profile cross-correlation matrix of BigBrain to identify principle axes of cytoarchitectural differentiation (Paquola et al., 2019) (**Figure 1D**). Here, we replicated the approach with updated staining intensity profiles. Bearing in mind the high-dimensional matrix manipulation necessary for this procedure, we first decimated the BigBrain mesh from 327,684 to ~10,000 vertices. Mesh decimation involves selection of a subset of vertices that preserve the overall shape of the surface followed by retriangulation of the faces with only the selected vertices. We assigned non-selected vertices to the nearest selected vertex, based on shortest path on the mesh (ties were solved by shortest Euclidean distance). In this manner, all 327,684 vertices belong to one of ~10,000 parcels. Derivation of the cytoarchitectural gradients involved (i) averaging staining intensity profiles within each parcel, (ii) pair-wise correlation of parcel-average staining intensity profiles (controlling for the global-average staining intensity profile), (iii) transformation to a normalised angle matrix, and iv) diffusion map embedding of this matrix. Each eigenvector captures an axis of cytoarchitectural variation and is accompanied by an eigenvalue that approximates the variance explained by that eigenvector. Here, the first two eigenvectors explain approximately 42% and 35% of variance, respectively, and describe anterior-posterior and sensory-fugal axes (further details in **Tutorial 2**).

### Morphometric models in BigBrain

The high resolution of BigBrain allows for precise segmentation of anatomical structures. Manual segmentations of the putamen, caudate nucleus, globus pallidus pars externa, globus pallidus pars interna, nucleus accumbens, amygdala, thalamus, red nucleus, substantia nigra, subthalamic nucleus and the hippocampus are available on Open Science Framework (https://osf.io/xkqb3/). Extending upon whole-structure segmentation, a recent study (DeKraker et al., 2019) used anatomical landmarks to create an internal coordinate system of the hippocampus. The approach involved solving Laplace’s equation under three sets of boundary conditions: anterior-posterior, proximal-distal (relative to the subiculum), and inner-outer (DeKraker et al., 2018). Subsequently, the hippocampus can be “unfolded”, allowing examination of histological and morphometric features in a topologically continuous space (**Figure 1E**), in line with other surface-based studies of the hippocampus (Bernhardt et al., 2016; Caldairou et al., 2016; Kim et al., 2014; Vos de Wael et al., 2018). Furthermore, this 3D coordinate system enabled the creation of a continuous surface model of the mesiotemporal cortex (Paquola et al., 2020b). The hippocampus is typically excluded from cortical surface models due to its complex folding and unusual cytoarchitectural makeup, with Cornu Ammonis subfields being allocortical and the dentate gyrus an interlocked terminus. Using the proximal-distal axis of the hippocampus, we were able to bridge the isocortical and hippocampal surface models recapitulating the smooth confluence of cortical types in the mesiotemporal lobe (**Figure 1F)**. The continuous surface model, defined by a pial/inner surface and a white/outer surface, can also be used to initialise equivolumetric surface constructions (Waehnert et al., 2014; Wagstyl et al., 2018b). We generated staining intensity profiles using 40μm resolution blocks of BigBrain across the cortical confluence, which are released in BigBrainWarp with the matching surface model.

### BigBrain-MRI transformations

BigBrain-MRI integration is pillared upon transformations between spaces. Spatial registration already exists as a fundamental component of most neuroimaging pipelines. As such, extensive research has focused on the creation of standard spaces, such as ICBM-MNI152 (Fonov et al., 2011b, 2009) and FreeSurfer’s fsaverage (Fischl et al., 1999). Multiple studies have demonstrated the continuous enhancement of registration techniques over the years (Collins and Evans, 1997; Klein et al., 2009; Xiao et al., 2019). Registration of BigBrain to MRI templates involves additional challenges, however, including histological artefacts, differences in intensity contrasts and inter-individual variability.

For the initial BigBrain release (Amunts et al., 2013), full BigBrain volumes were resampled to ICBM2009sym (a symmetric and non-linear MNI152 template) and MNI-ADNI (an older adult T1-weighted template) (Fonov et al., 2011a). Each resampling procedure involved a linear then a nonlinear transformation (available on ftp://bigbrain.loris.ca/BigBrainRelease.2015/). BigBrain volumes resampled to ICBM2009sym are commonly referred to as BigBrainSym. We continue to use this nomenclature in BigBrainWarp. A prior study (Xiao et al., 2019) was able to further improve the accuracy of the transformation for subcortical structures and the hippocampus using a two-stage multi-contrast registration procedure. The first stage involved nonlinear registration of BigBrainSym to a PD25 T1-T2* fusion atlas (Xiao et al., 2017, 2015), using manual segmentations of the basal ganglia, red nucleus, thalamus, amygdala, and hippocampus as additional shape priors. Notably, the PD25 T1-T2* fusion contrast is more similar to the BigBrainSym intensity contrast than a T1-weighted image, such as the commonly used ICBM2009sym template. The second stage involved nonlinear registration of PD25 to ICBM2009sym and ICBM2009asym using multiple contrasts. The authors have shared the deformation matrices on Open Science Framework (https://osf.io/xkqb3/). The accuracy of the transformations was evaluated relative to anatomical fiducials (Lau et al., 2019) and regional segmentations. The two-stage procedure resulted in 0.86-0.97 DICE coefficients for manual segmentations, improving upon direct overlap of BigBrainSym with ICBM2009sym (0.55-0.91 DICE). Anatomical fiducials alignment incurred 1.77±1.25mm errors, on par with direct overlap of BigBrainSym with ICBM2009sym (1.83±1.47mm). In line with this work, BigBrainWarp enables evaluation of novel deformation fields using anatomical fiducials (Lau et al., 2019) and region segmentations (**evaluate_warps.sh**).

The unique morphology of BigBrain also presents challenges for surface-based transformations. Idiosyncratic gyrification of certain regions of BigBrain, especially the anterior cingulate, cause misregistration (Lewis et al., 2020). To overcome this issue, ongoing work leverages multimodal surface matching [MSM; (Robinson et al., 2018, 2014)] to optimise surface transformation from BigBrain to standard surface templates. This procedure improves accuracy and minimises distortion of transformed cortical maps, almost on par with *in vivo* MRI transformations (Lewis et al., 2020).

### Compiling BigBrainWarp

For BigBrainWarp, we wrote a modular set of wrapper scripts to map between common BigBrain and MRI spaces (**Figure 2**). The package automatically pulls state-of-the-art deformation matrices, then applies the transformation to novel data. While applying these various transformations involve different tools (*e.g.:* minc-tools, FSL, HCP-workbench), BigBrainWarp wraps these functions into a single bash script (see **Table 2** for functionality), reducing onus on the user to have experience in each software package. Furthermore, containerisation of the BigBrainWarp via Docker allows users to interact with the scripts without installing dependencies. This procedure ensures flexibility with ongoing developments in the field and simplifies procedures for new users.

**Figure 2:**
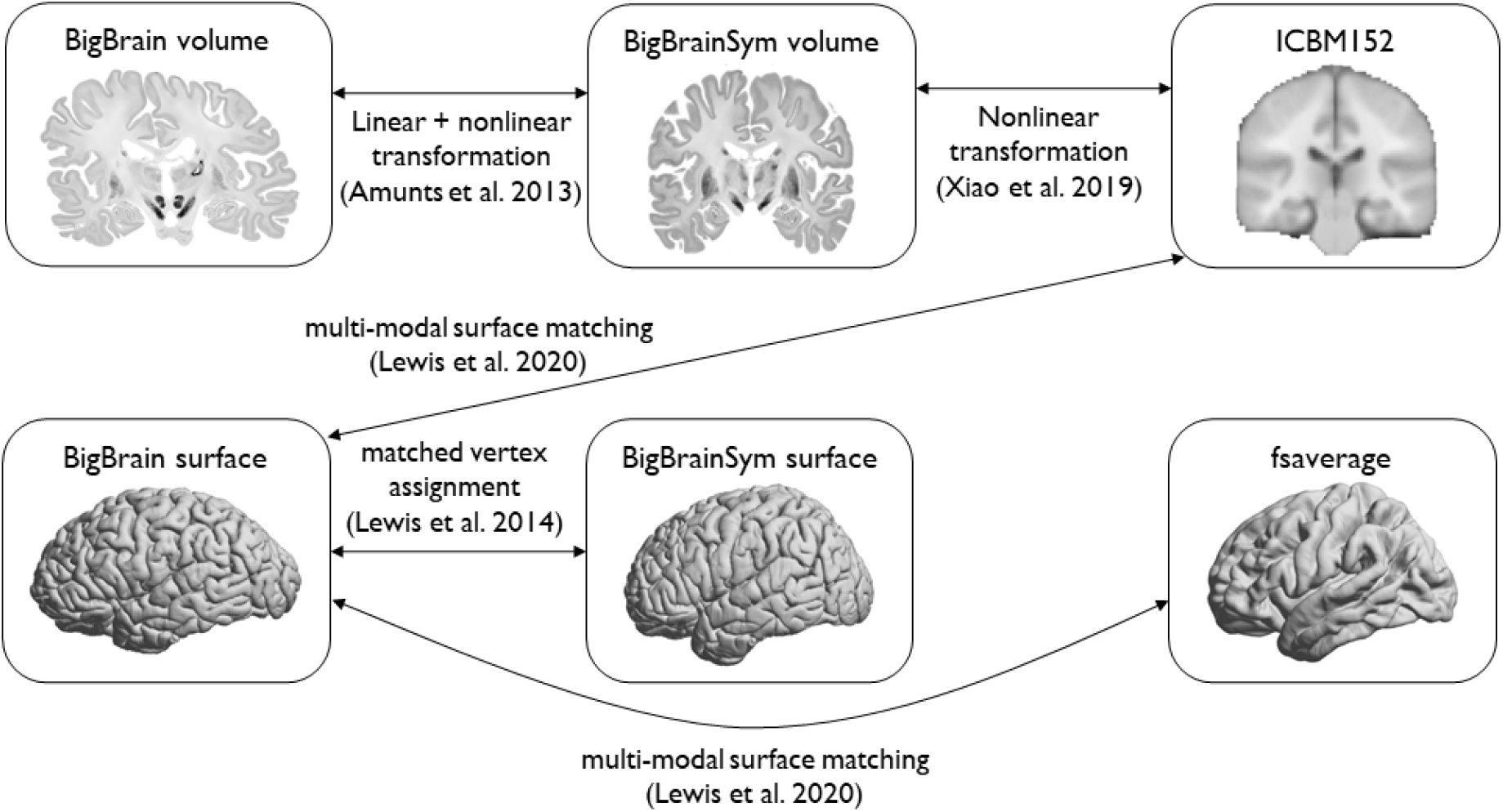
Overview of spaces and transformations included within BigBrainWarp.

**Table 2:**
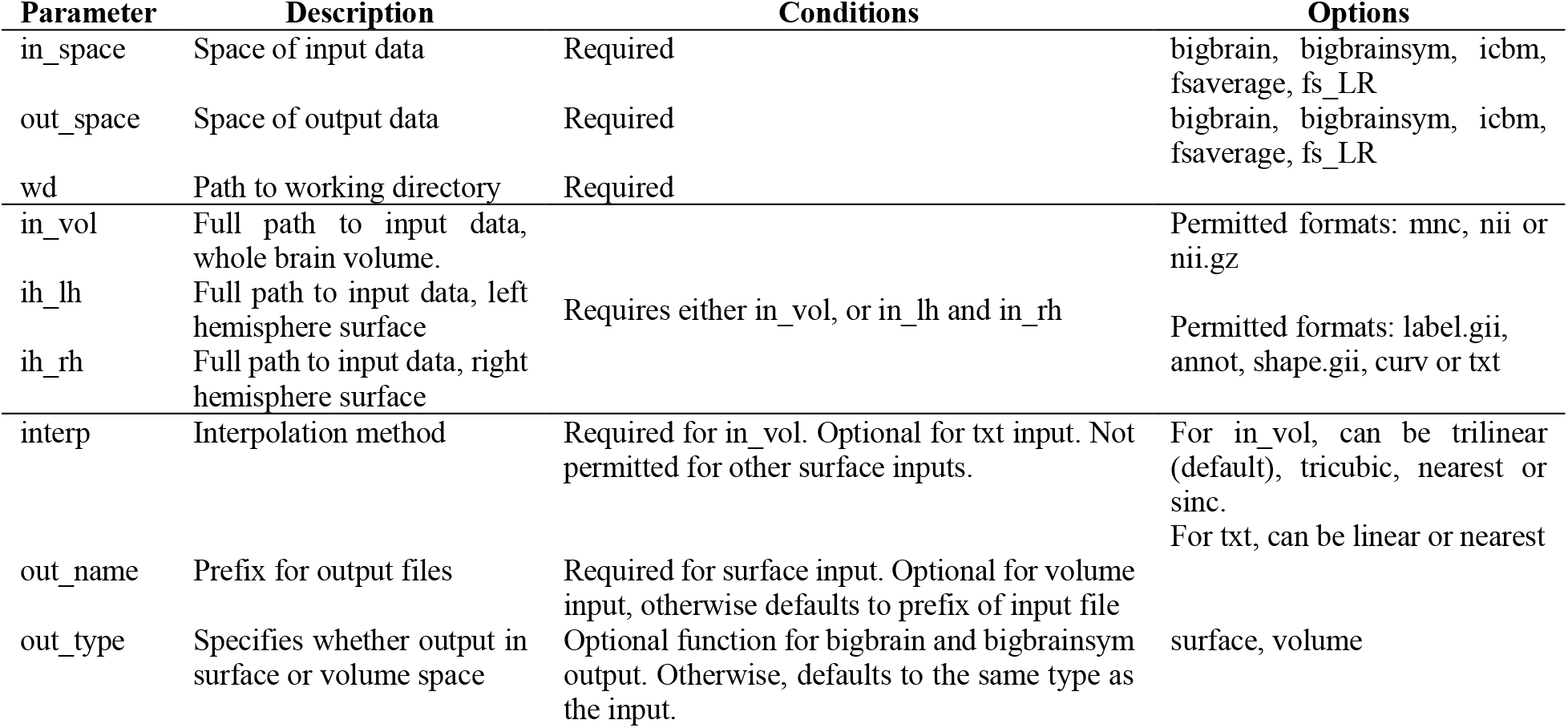
Input parameters for the bigbrainwarp function

We used BigBrainWarp to map histological gradients to fsaverage, fs_LR and ICBM152. For the initial release of BigBrainWarp, we selected a multi-scale imaging dataset (MICs), which contains group-level imaging features on standard surface templates from 50 healthy adults. In particular, we adopted cortical gradients derived from qT1 mapping and resting-state functional connectivity. We used BigBrainWarp to transform microstructural and functional gradients, as well as intrinsic functional communities (Yeo et al., 2011), to the BigBrain surface. The current contents of the toolbox are shown in **Table 3**.

**Table 3:**
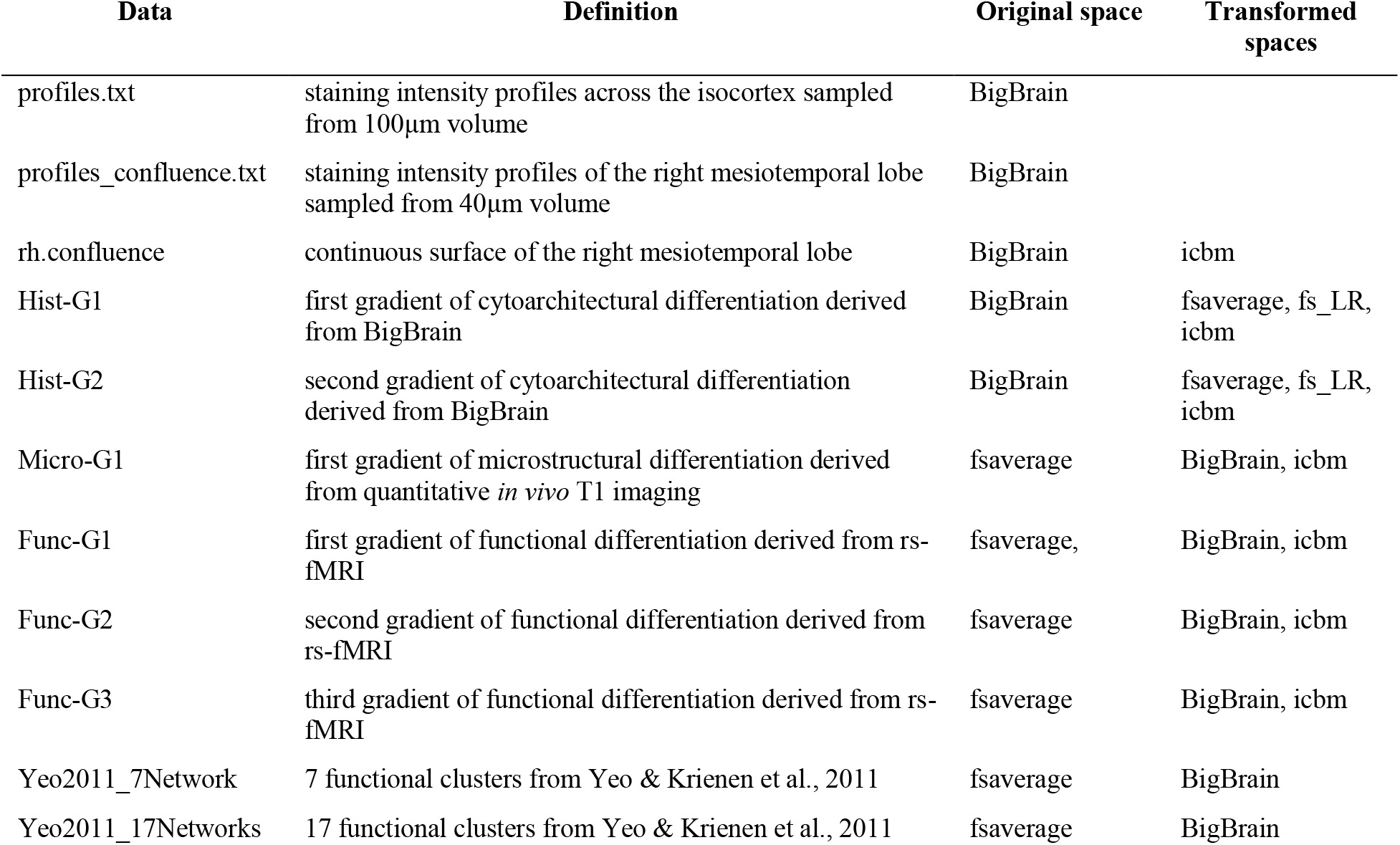
BigBrainWarp contents.

| Data | Definition | Original space | Transformed spaces |
| --- | --- | --- | --- |
| profiles.txt | staining intensity profiles across the isocortex sampled from 100μm volume | BigBrain | |
| profiles_confluence.txt | staining intensity profiles of the right mesiotemporal lobe sampled from 40μm volume | BigBrain | |
| rh.confluence | continuous surface of the right mesiotemporal lobe | BigBrain | icbm |
| Hist-G1 | first gradient of cytoarchitectural differentiation derived from BigBrain | BigBrain | fsaverage, fs_LR, icbm |
| Hist-G2 | second gradient of cytoarchitectural differentiation derived from BigBrain | BigBrain | fsaverage, fs_LR, icbm |
| Micro-G1 | first gradient of microstructural differentiation derived from quantitative *in vivo* T1 imaging | fsaverage | BigBrain, icbm |
| Func-G1 | first gradient of functional differentiation derived from rs-fMRI | fsaverage, | BigBrain, icbm |
| Func-G2 | second gradient of functional differentiation derived from rs-fMRI | fsaverage | BigBrain, icbm |
| Func-G3 | third gradient of functional differentiation derived from rs-fMRI | fsaverage | BigBrain, icbm |
| Yeo2011_7Network | 7 functional clusters from Yeo & Krienen et al., 2011 | fsaverage | BigBrain |
| Yeo2011_17Networks | 17 functional clusters from Yeo & Krienen et al., 2011 | fsaverage | BigBrain |

## 3. Results

The BigBrainWarp toolbox supports a range of integrative BigBrain-MRI analyses. The following tutorials outline three BigBrain-MRI analyses with unique types of transformations. Neither the forms nor the motivations are exhaustive but illustrate applications. Code for each tutorial is available in the BigBrainWarp toolbox.

### Tutorial 1: BigBrain → ICBM2009sym MNI152 space

#### Motivation

Despite MRI acquisitions at high and ultra-high fields reaching submillimeter resolutions with ongoing technical advances, certain brain structures (*e.g.,* subthalamic nucleus) and subregions (*e.g.,* hippocampal Cornu ammonis subfields) remain difficult to identify (Kulaga-Yoskovitz et al., 2015; Wisse et al., 2017; Yushkevich et al., 2015). BigBrain can be used to label such regions, then the atlas labels can be transformed to a standard imaging space for further investigation. In particular, this approach can support exploration of the functional architecture of histologically-defined regions of interest.

#### Approach

(i) Create volumetric label in BigBrain space. (ii) Perform nonlinear transformation to ICBM2009sym space using BigBrainWarp. (iii) Transform individual resting-state functional MRI data to ICBM2009sym MNI152 space. (iv) Sample timeseries from labelled voxels in this standard space.

#### Example

The mesiotemporal lobe plays important roles in multiple cognitive processes (Moscovitch et al., 2005; Squire et al., 2004; Vos de Wael et al., 2018) and is affected by multiple neurological and neuropsychiatric conditions (Ball et al., 1985; Bernhardt et al., 2016, 2015; Calabresi et al., 2013). Increasing research suggests that this region shows complex subregional structural and functional organization. Here, we illustrate how we track resting-state functional connectivity changes along the latero-medial axis of the mesiotemporal lobe, from parahippocampal isocortex towards hippocampal allocortex. For further details and additional motivation, please see (Paquola et al., 2020a): (i) Our volumetric label represents the iso-to-allocortical axis of the mesiotemporal lobe. We constructed this axis by joining the isocortical (Lewis et al., 2014) and hippocampal (DeKraker et al., 2019) surface meshes in BigBrain histological space, calculated the distance of each vertex in the new surface model to the intersection of isocortical and hippocampal meshes (**Figure 3A**). Next, we labelled voxels in BigBrain histological space, according to the position of the iso-to-allocortical axis (**Figure 3Bii**). The iso-to-allocortical axis is ready-made in the BigBrainWarp toolbox. (ii) We transform the volume from the BigBrain histological space to ICBM2009sym (**Figure 3Biii**). (iii) For each participant, in this case 50 healthy adults from the MICs dataset, we construct an individualised transformation from ICBM2009sym to native functional space, based on the inverse of the within-subject co-registration to the native T1-weighted imaging concatenated to the nonlinear between-subject registration to ICBM2009sym. (iv) For each participant, BOLD timeseries are extracted from non-zero voxels of the transformed iso-to-allocortical axis, which are classified as grey matter (>50% probability) and collated in a 3D matrix (voxel × time × subject). Then, we sort and analyse this matrix using the voxel-wise values of the iso-to-allocortical axis. For instance, product-moment correlations of strength of resting state functional connectivity with iso-to-allocortical axis indicates how functional connectivity varies along the histological axis for different areas of the isocortex (**Figure 3C**).

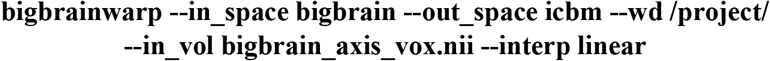

**Figure 3:**
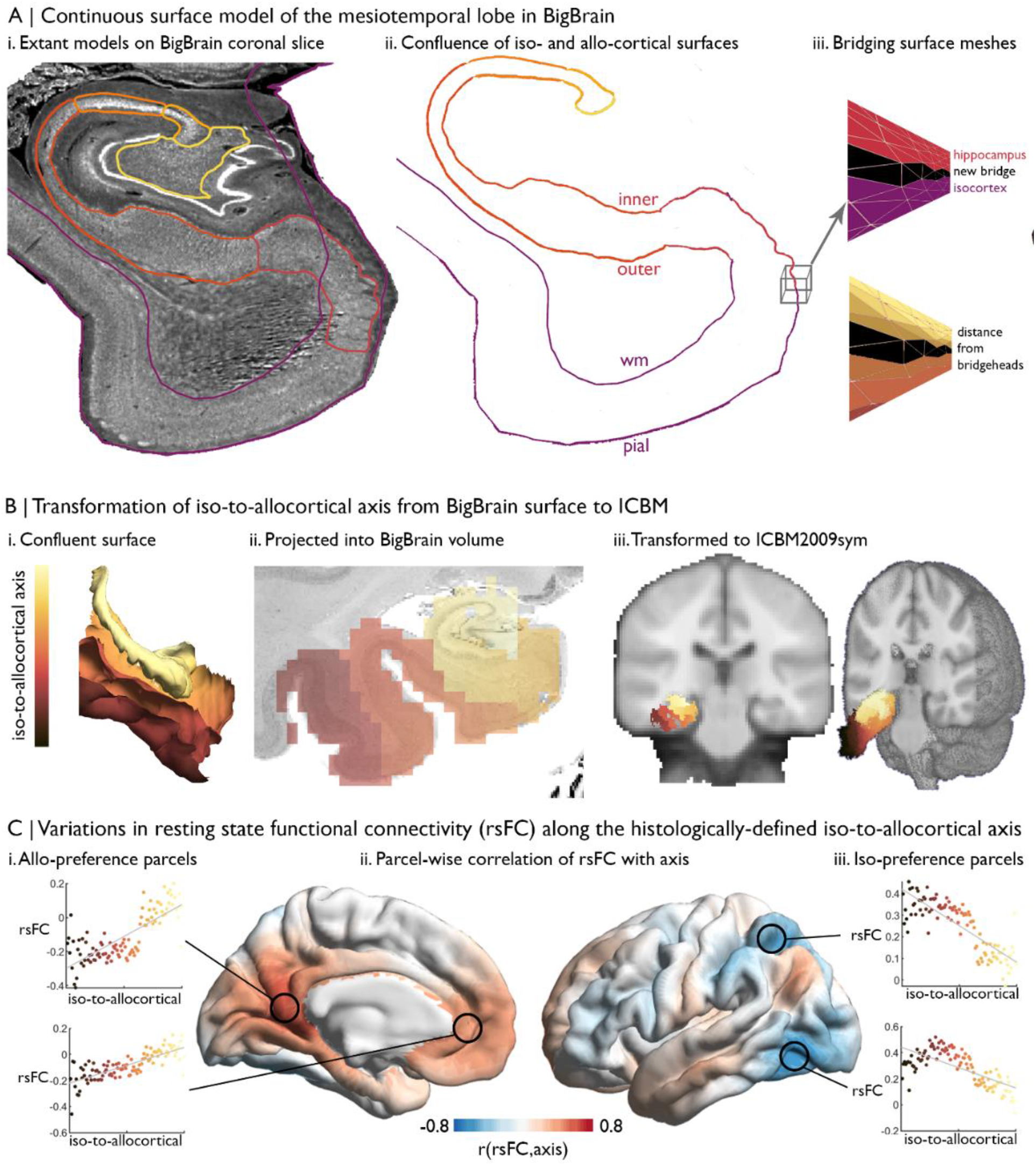
Intrinsic functional connectivity of the iso-to-allocortical axis of the mesiotemporal lobe. **A) i.** BigBrain surface models of the isocortex and hippocampal subfields are projected on a 40 μm resolution coronal slice of BigBrain. **ii-iii**. The continuous surface model bridges the inner hippocampal vertices (minimum value on inner-outer axis) with pial mesiotemporal vertices (entorhinal, parahippocampal or fusiform cortex). Vertices at the medial aspect of the subiculum were identified as bridgeheads and used to bridge between the two surface constructions. Geodesic distance from the nearest bridgehead was used as the iso-to-allocortical axis. **B)** Iso-to-allocortical axis values were projected from the surface into the BigBrain volume, then transformed to ICBM2009sym MNI152 space using BigBrainWarp. **C)** Intrinsic functional connectivity was calculated between each voxel of the iso-to-allocortical axis and 1000 isocortical parcels, using rs-fMRI images nonlinearly registered to ICBM2009sym. For each parcel, we calculated the product-moment correlation of rsFC strength with iso-to-allocortical axis position.

### Tutorial 2: BigBrain → fsaverage

#### Motivation

*In vivo* brain imaging reveals regionally variable effects of many demographic and clinical factors on brain structure and function. For example, prior studies studying lifespan processes presented spatially variable patterns of cortical atrophy with advancing age, together with increased deposition of pathological aggregates, such as amyloid beta (Bilgel et al., 2018; Jansen et al., 2015; Knopman et al., 2018; Rodrigue et al., 2012; Sperling et al., 2011). Histological data provides a window into the cytoarchitectural features that align with imaging-derived phenotypes and that, in this instance, may predispose an area to specific aging related processes. Essentially, we can evaluate whether regions with a certain cytoarchitecture overlap with those showing more marked aging effects. Furthermore, large-scale cytoarchitectural gradients can provide a unified framework to describe topographies, simplifying and standardising the reporting of imaging-derived phenotypes.

#### Approach

(i) Construct histological gradients using BigBrain and (ii) transform to standard neuroimaging surface template using BigBrainWarp. (iii) Plot the imaging-derived map against each histological gradient to understand the algebraic form of the relationship. Note, if imaging features are volumetric, one may use registration fusion to resample the data from ICBM2009sym to fsaverage (Wu et al., 2018). (iv) Fit a statistical model to evaluate the relationship between the cytoarchitectural gradients and the imaging-derived map. For research questions with a more restricted region of interest, the cytoarchitectural gradient could be reconstructed within that field of view and the same procedure could be utilised. The optimal number of cytoarchitectural gradients should be evaluated.

#### Example

Cytoarchitectural correlates of age-related increases in amyloid beta (Aβ) deposition in a healthy lifespan cohort (Lowe et al., 2019; Park, 2018). (i) and (ii) are pre-computed in BigBrainWarp (**Figure 4A**) using

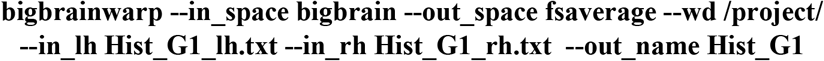

**Figure 4:**
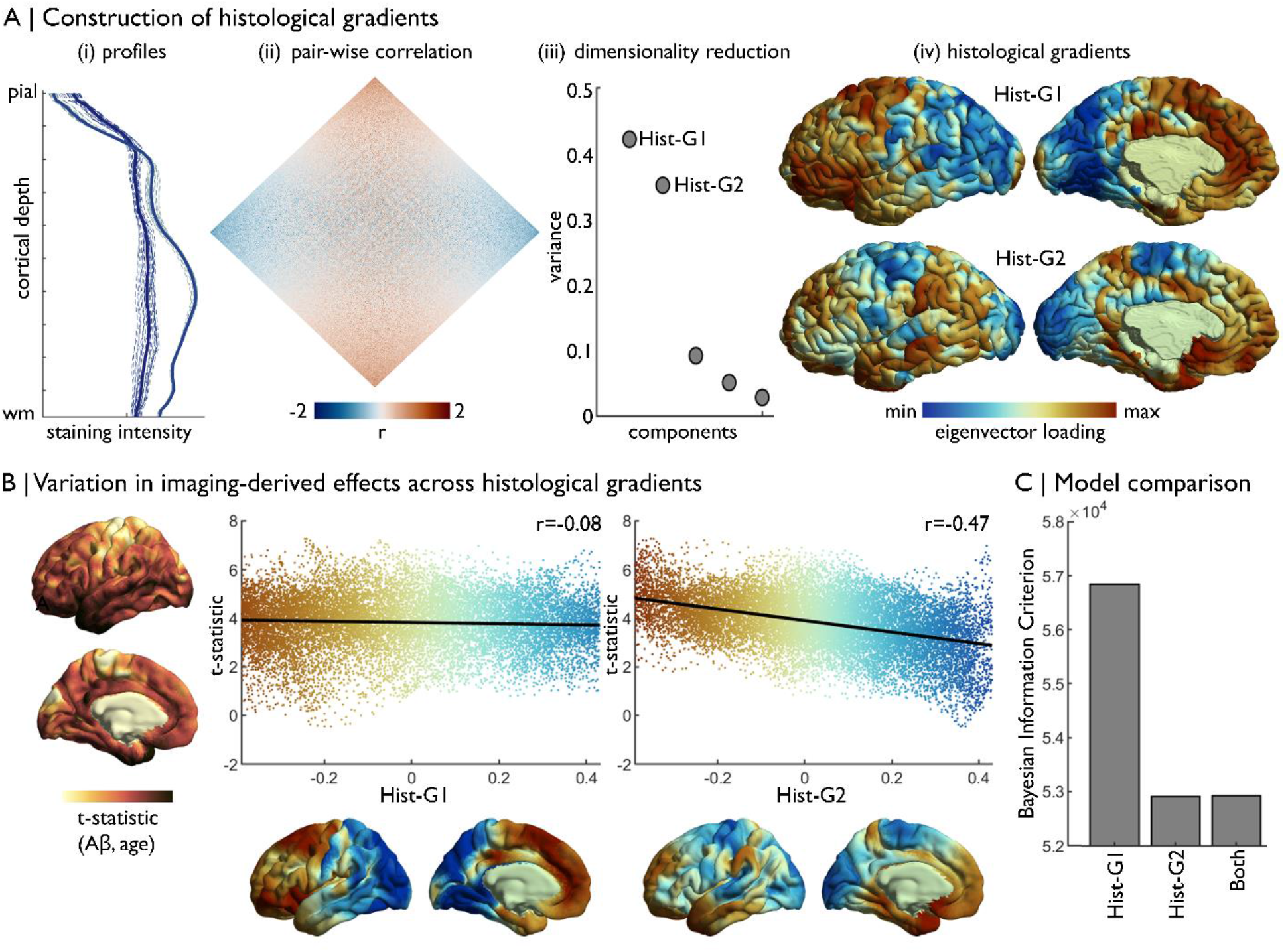
Concordance of imaging-derived effects with histological gradients. **A)** Four stages of histological gradient construction. (i) Vertex-wise staining intensity profiles (dotted lines) are averaged within parcels (solid lines). Colours represent different parcels. (ii) Pair-wise partial correlation of parcel-average staining intensity profiles produces a cortex-wide matrix of cytoarchitectural similarity. (iii) The correlation matrix is subjected to dimensionality reduction, in this case diffusion map embedding, to extract the principle axes of cytoarchitectural variation. (iv) The principle components capture histological gradients (Hist-G) and are projected onto the BigBrain cortical surface for inspection. **B)** The t-statistic cortical map illustrates regional variations in the effect of age on Aβ deposition (Lowe et al., 2019), which was calculated vertex-wise on fsaverage5. To allow comparison, histological gradients were transformed to fsaverage5 using BigBrainWarp. Scatterplots show the association of the t-statistic map with the histological gradients. **C)** Bar plot shows the Bayesian Information Criterion of univariate and multivariate regression models, using histological gradients to prediction regional variation in effect of age on Aβ deposition. The univariate Hist-G2 regression had the lowest Bayesian Information Criterion, representing the optimal model of those tested.

For this analysis, we used a 6mm FWHM smoothing kernel to approximately match the smoothing kernel of the resting state fMRI data. (iii) We previously estimated the association of age with amyloid deposition across the cortical surface by combining positron emission tomography with MRI data in 102 adults (30-89 years), and assessed correspondence to functional connectivity gradients (Lowe et al., 2019). Here, we plot the vertex-wise t-statistics against Hist-G1 and Hist-G2 (**Figure 4B**) (iv) We determine the optimal model via the Bayesian Information Criterion in univariate and multivariate regressions between the t-statistics and histological gradients (**Figure 4C**). The optimal model included only Hist-G2, indicating that Aβ preferentially accumulates towards the more agranular anchor of the sensory-fugal gradient.

### Tutorial 3: fsaverage/ICBM2009sym → BigBrain

#### Motivation

A core aim of fMRI research is to map functional specialisation in the brain (Bassett et al., 2008; Eickhoff et al., 2018; Gordon et al., 2017; Raichle, 2015; Shine et al., 2019; Yeo et al., 2011). On the one hand, this work follows a long legacy of defining cortical areas, and on the other hand, it extends beyond the possibilities of *post mortem* research by capturing patterns of coordinated activity. For instance, clustering resting state fMRI connectivity reveals a robust set of intrinsic functional networks (Beckmann and Smith, 2004; Gordon et al., 2017; Yeo et al., 2011). Nonetheless, there exists a gap in the literature between these well-characterised functional networks and their cytoarchitecture. BigBrain offers the opportunity to characterise and evaluate differences of cytoarchitecture for functionally defined atlases.

#### Approach

(i) Transform functionally-defined regions from a standard neuroimaging surface template to the BigBrain surface. Note, if the functional-defined regions are volumetric, one may use registration fusion to resample the data from ICBM2009sym to fsaverage (Wu et al., 2018). (ii) Compile staining intensity profiles by functional class. (iii) Assess discriminability of functional classes by staining intensity profiles.

#### Example

Cytoarchitectural differences of intrinsic functional networks. (i) Transform the 17-network functional atlas (Yeo et al., 2011) to the BigBrain surface.

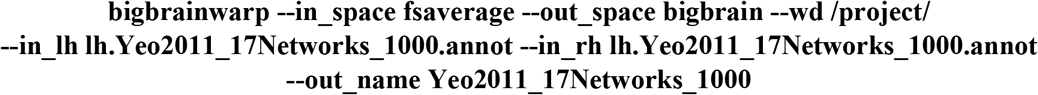

(ii) Stratify staining intensity profiles by network (**Figure 5A**). (iii) Parameterise staining intensity profiles by the central moments and assess variation across functional networks (**Figure 5B**). For example, the mean and skewness illustrate distinct patterns of cytoarchitectural differentiation across the functional networks. Visual networks have the highest mean and lowest skewness. Somatomotor, dorsal attention and fronto-parietal networks contain most variable mean and skewness values. Ventral attention, limbic and fronto-parietal networks harbour the lowest mean and highest skewness, whereas the default mode networks occupy an intermediary position. Notably, all the networks exhibit broad distribution of the moments, signifying substantial cytoarchitectural heterogeneity, as well as overlapping values. To quantify discriminability of functional networks by cytoarchitecture, we can attempt to classify the functional networks using the central moments. For this example, we z-standardised the central moments and split the vertices into five folds, each with an equal representation of the 17 functional networks. Then, we trained a one vs one linear support vector classification on 50% of each fold and tested the model on the remaining 50% of that fold. Functional networks were equally stratified across training and testing. Finally, for each fold, we generated a confusion matrix, showing the accurate predictions on the diagonal and the incorrect classification off the diagonal. Predictive ability provides insight into distinctiveness and homogeneity of functional networks. Visual networks harbour distinctive cytoarchitecture, reflected by relatively high accuracy and few incorrect predictions. Ventral attention, limbic and temporoparietal networks are relatively homogenous in cytoarchitecture, likely related to their restricted spatial distribution. The predictive accuracy did not appear to be negatively impacted by minor misalignments of the atlas, as the predictive accuracy was similar when excluding vertices within approximately 6mm of the network boundaries (accuracy mean±SD (%), original=12.4±15.4, excluding boundaries=12.1±13.3).

**Figure 5:**
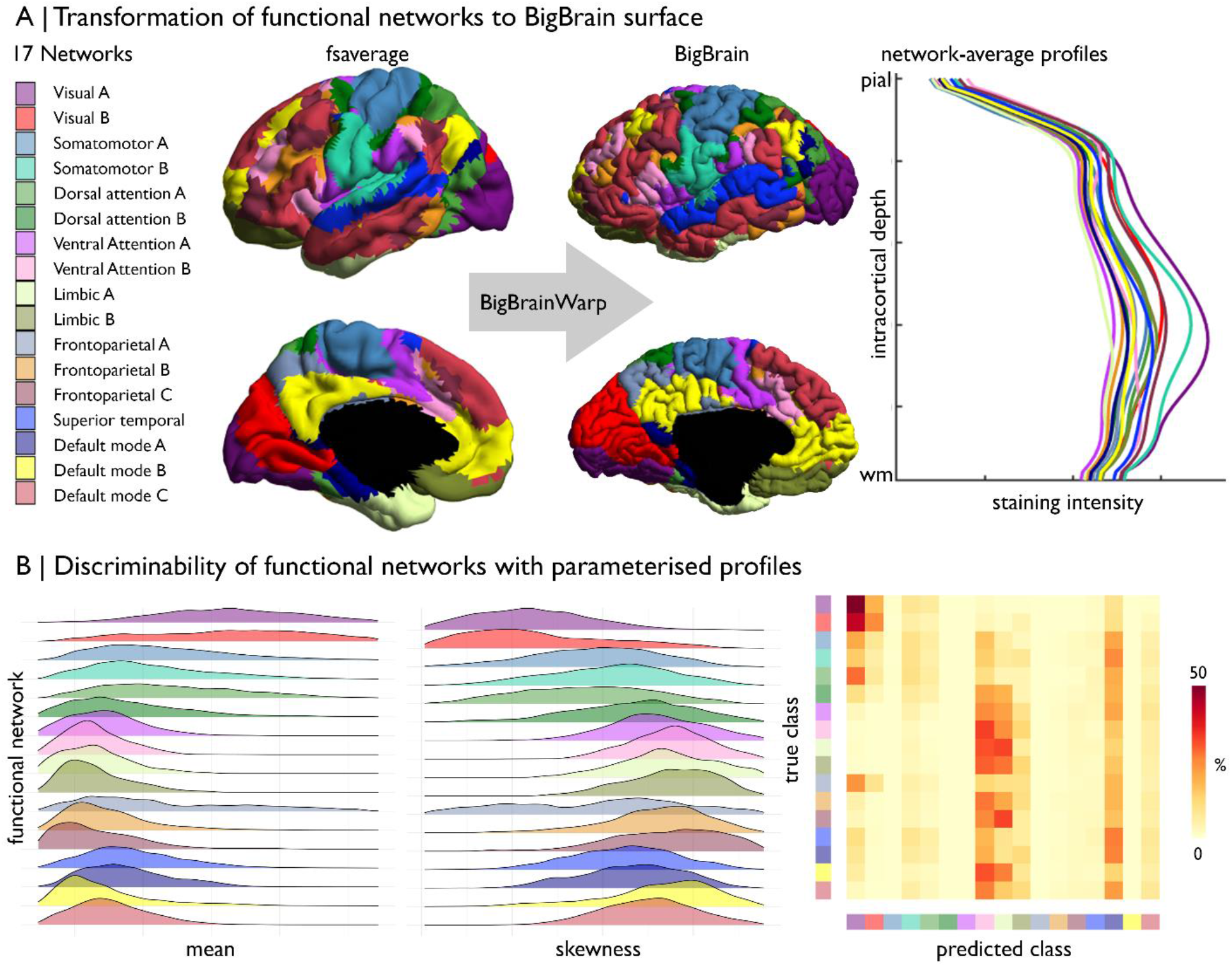
Prediction of functional network by cytoarchitecture. **A)** Surface-based transformation of 17-network functional atlas to the BigBrain surface, operationalised with BigBrainWarp, allows staining intensity profiles to be stratified by functional network. **B)** Ridgeplots show the moment-based parameterisation of staining intensity profiles within each functional network. The confusion matrix illustrates the outcome of mutli-class classification of the functional networks, using the central moment of the staining intensity profiles.

## 4. Discussion

Beyond cartography, a major aim of neuroanatomical research has been to understand the functioning of the human brain. Throughout the 20^th^ century, cytoarchitectural studies were instrumental in demonstrating functional specialisation across the cortex, as well as the uniqueness of the human brain amongst mammals (Brodmann, 1909; Campbell et al., 1905; Sanides, 1962; Smith, 1907; Vogt and Vogt, 1919; Vogt, 1911). Fine-grained anatomical resolution maintains an important role in understanding brain function in the modern era, helping to bridge between microcircuit organisation and macroscale findings obtained with *in vivo* neuroimaging. BigBrain is the first ultra-high-resolution 3D histological dataset that can be readily integrated with *in vivo* neuroimaging. In this report, we presented BigBrainWarp, a simple and accessible toolbox comprising histological data, previously developed transformation functions between BigBrain and standard imaging spaces, and ready-to-use transformed cortical maps. The toolbox is containerised to eliminate software dependencies and to ensure reproducibility. An expandable documentation is available, alongside several tutorials, at http://bigbrainwarp.readthedocs.io.

Multimodal registrations are core to integrating BigBrain with *in vivo* neuroimaging data. Identifying optimal solutions is more difficult than intra- and inter-subject co-registrations of neuroimaging data, owing to histological artefacts, differences in intensity contrasts and morphological distortions. These challenges have been addressed by recent studies, which improved integration of BigBrain with standardised MRI spaces. An automated repair algorithm was specially devised for BigBrain, which involved nonlinear alignment of neighbouring sections, intensity normalisation, outlier detection using block averaging then artefact repair using the block averages (Lepage et al., 2010; Lewis et al., 2014). Following initial transformation of BigBrain to ICBM2009b, which was part of the initial BigBrain release (Amunts et al. 2013), a recent study optimised subcortical registrations by generating a T1-T2* fusion contrast that is more similar to the BigBrain intensity contrast than a T1-weighted image (Xiao et al., 2019). Additionally, that study involved manual segmentation of subcortical nuclei to use as shape priors in the registration, which benefits the alignment of subcortical structures between BigBrain and standard neuroimaging templates. Finally, inspired by advances in the alignment of surface-based MRI data (Robinson et al., 2018, 2014), the BigBrain team has recently developed a multi-modal surface matching pipeline for BigBrain that involved re-tessellation of the BigBrain surface at a higher resolution, followed by alignment to standard surface templates using coordinate, sulcal depth and curvature maps (Lewis et al., 2020). The procedure significantly improves upon previous techniques, resulting in geometric distortions comparable to those seen for registrations between neuroimaging datasets of different individuals (Lewis et al., 2020).

Practically, 3D histological models provide an unrivalled level of precision, and provide novel opportunities to cross-validate and contextualise findings from human neuroimaging. BigBrainWarp is particularly well-suited for investigations on the fundamental relationships between cytoarchitecture and function, which remains an elusive aspect of brain organisation. Our tutorials illustrate and deconstruct a range of use cases of BigBrain-MRI integration. In tutorial 1, we show how BigBrain can be used to initialise region of interest analyses, such as mapping resting state functional connectivity along the iso-to-allocortical axis (Paquola et al., 2020b), enabling precise delineation of regions that are difficult to identify with *in vivo* imaging and functional interrogation of histological axes. In tutorial 2, we show how cytoarchitectural gradients can help to characterise large-scale cortical patterns, such as the association of aging with Aβ deposition (Lowe et al., 2019). This approach complements the tradition of reporting the cortical areas of significant clusters by offering a simplified topographical description of the spatial pattern. Furthermore, by comparing predictive power of various cytoarchitectural gradients, we may build towards hypotheses on the relationship between microcircuit properties and demographic or clinical factors. In tutorial 3, we discuss more specific histological features, namely moment-based parameterisation of staining intensity profiles (Schleicher et al., 1999; Zilles et al., 2002). These features depict the vast cytoarchitectural heterogeneity of the cortex and enable evaluation of homogeneity within imaging-based parcellations, for example macroscale functional communities (Yeo et al., 2011). Together, these tutorials showcase how we can easily and robustly use BigBrain with BigBrainWarp to deepen our understanding of the human brain.

Despite all its promises, the singular nature of BigBrain currently prohibits replication and does not capture important inter-individual variation at the scale of histology. Fortunately, the BigBrain teams are working on new histology-based 3D models in the context of the HIBALL project (https://bigbrainproject.org/hiball.html). System neuroscience has dramatically benefitted from the availability of open resources (Di Martino et al., 2014; Milham et al., 2018; Poldrack et al., 2017; Van Essen et al., 2013). This path, together with ongoing refinements in multimodal data integration and efforts to make tools accessible, promises to further advance multi-scale neuroscience in the years to come.

## Acknowledgements

The project was conducted as part of the Helmholtz International BigBrain Analytics Learning Laboratory (HIBALL), an international initiative funded by Helmholtz Association & Healthy Brains for Healthy Lives. Casey Paquola was funded through the Fonds de la Recherche du Quebec – Santé (FRQ-S). Boris Bernhardt acknowledges research support from the National Science and Engineering Research Council of Canada (NSERC Discovery-1304413), the Canadian Institutes of Health Research (CIHR FDN-154298, PRJ-174995), SickKids Foundation (NI17-039), Azrieli Center for Autism Research (ACAR-TACC), BrainCanada (Azrieli Future Leaders), the Tier-2 Canada Research Chairs program and FRQ-S. Jessica Royer received support from a Canadian Institute of Health Research (CIHR) Fellowship. Ali Khan acknowledges research support from CIHR Project Grant #366062, NSERC Discovery Grant #6639, and the Canada First Research Excellence Fund.

## Supplementary Material

**Supplementary Figure 1:**
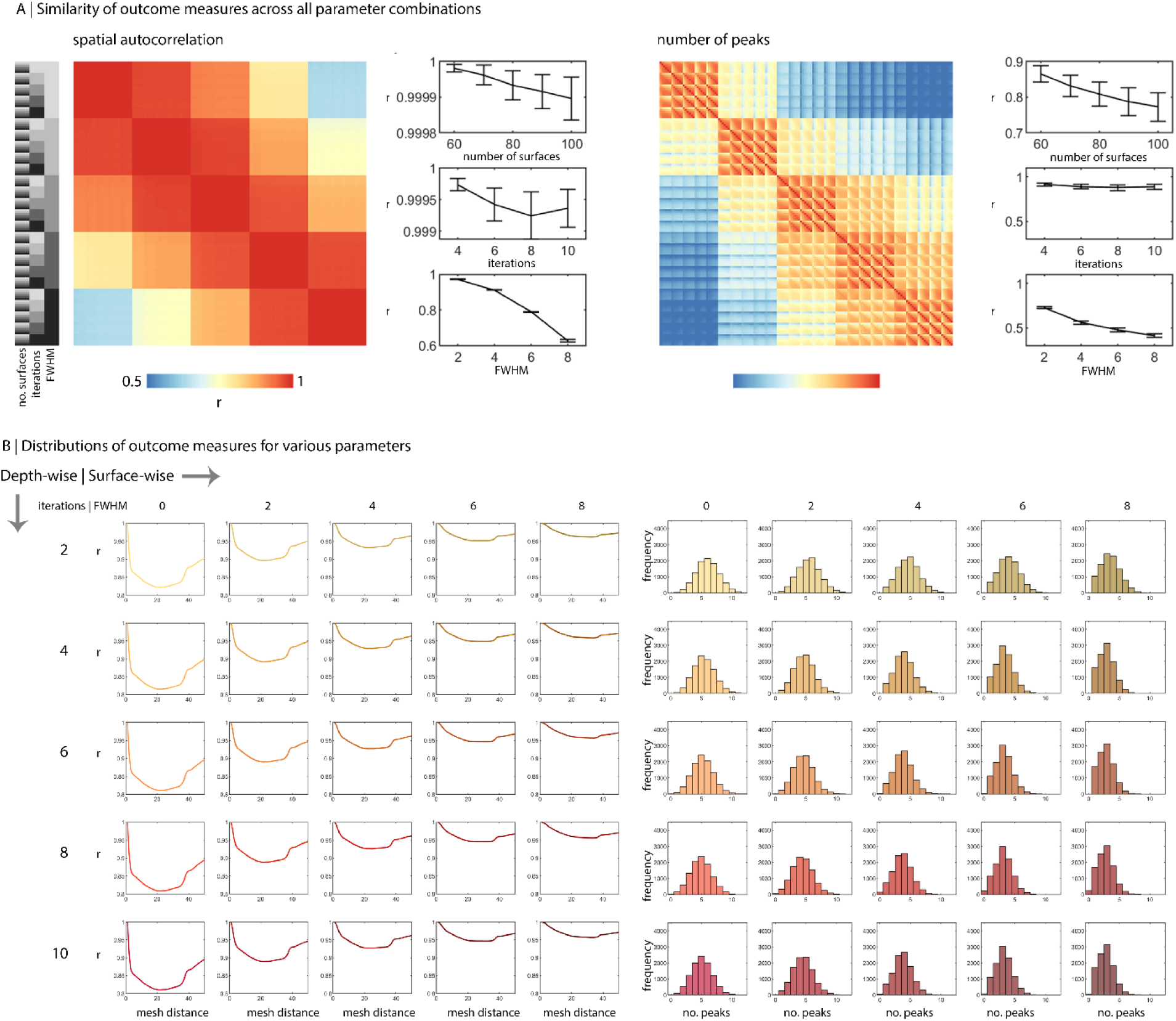
Evaluation of sampling parameters for staining intensity profiles. **A)** Matrices show the similarity (r) of spatial autocorrelation and number of peaks between parameter combinations. On the far left, grey bars show the parameter combination for each row of the matrix. Errorbar plots show the mean and SD of the correlation across a given parameter, while the other two parameters are consistent. The correlations are shown with respect to the lowest of each parameter (50 surfaces, 2 iterations and 0 FWHM). **B)** For varying degrees of depth-wise (rows) and surface-wise (columns) smoothing, line plots show spatial autocorrelation and histograms show number of peaks.

